# Spinal cord population activity lacks rotational dynamics during an alternating isometric task in macaque

**DOI:** 10.64898/2026.06.29.735164

**Authors:** Omar Refy, Steve I. Perlmutter, Marc A. Maier, Wade S. Smith, Eberhard E. Fetz, Jens Bo Nielsen

## Abstract

Recent studies report rotational population dynamics in spinal cord neuronal activity during rhythmic movements, suggesting computational principles shared with motor cortex. Here we show that primate cervical spinal cord activity does not exhibit rotational dynamics during an alternating single-joint isometric wrist task, instead it displays low-dimensional alternating population patterns. Positive controls confirm presence of rotational structure in motor cortex activity during the same task, indicating distinct computational strategies across the motor axis. Cortical neurons with post-spike effects on motoneurons had activity with dynamics resembling cortical rather than spinal populations.

## Main

The firing rates of a population of n units can be represented by a point in n-dimensional state space, whose orthogonal axes represent the firing rate of each unit. During repetitive behavior the changes in firing rate generate closed loop trajectories of this point, whose features can be described by the principal components (PCs). The largest PCs capture the majority variance of the population dynamics during movement. If the firing rates of a population of units are phase-locked to different phases of the movement, such as in alternating torque production, and have their peak firing rates uniformly distributed throughout the movement, then the population variance can be adequately captured by two comparable PCs: the first PC captures activity of units firing with the two alternating directions of movement, and the second principal component captures activity of units firing in the transition between the alternating states. Such a pattern of activity, where most variance is captured by two orthogonal alternating PCs, is called rotational dynamics. The population variance of another pattern of activity, where most units fire with the two opposing directions of the movement, but few or none during the transition phase, can be captured by one dominant PC, and we refer to that pattern as alternating dynamics.

Motor cortex activity exhibits rotational population dynamics during reaching and other movements^1–3^, reflecting neural trajectory evolution in a low-dimensional state space. Whether similar dynamics operate throughout the motor system remains debated. In contrast with prior literature on half-center-based alternation^4–10^, recent studies identified rotational dynamics in lumbar spinal cord of turtles^11^, cats^12^, and mice^13^, suggesting that a rotational structure may be a general feature of motor circuits. However, these studies exclusively examined rhythmic multi-joint movements (locomotion, scratching), where sequential muscle activation across joints may have produced apparent rotational structure in state space independently of the underlying computational mechanism.

We tested whether spinal cord exhibits rotational dynamics during non-rhythmic but alternating single-joint muscle contractions by analyzing neural recordings from three male rhesus macaques performing isometric wrist torque tasks (Fig. 1A). Two monkeys (Monkeys B and W) provided cervical spinal cord recordings; a third monkey (Monkey D) provided motor cortex recordings as a control. Animals generated alternating flexion and extension torques with visual feedback following a ramp-and-hold profile. Extracellular single cell recordings were obtained from cervical spinal cord interneurons (C6–T1, ipsilateral to the torque-producing limb) and from contralateral primary motor cortex (hand area) using glass-coated tungsten electrodes^14–16^. Spike sorting yielded 100 well-isolated single units, 52 multi-unit clusters and 27 noisy clusters from Monkey B (152 total), and 88 single units with 81 multi-unit and 31 noisy clusters from Monkey W (169 total). From motor cortex, Monkey D yielded 159 single units, and 81 multi-unit and 94 noisy clusters (240 total) (Fig. 1F). Units were classified based on ISI violation rate, isolation distance, and waveform amplitude stability (see Methods). Histograms of task related neural activity were aligned with the torque signal.

**Figure 1.**
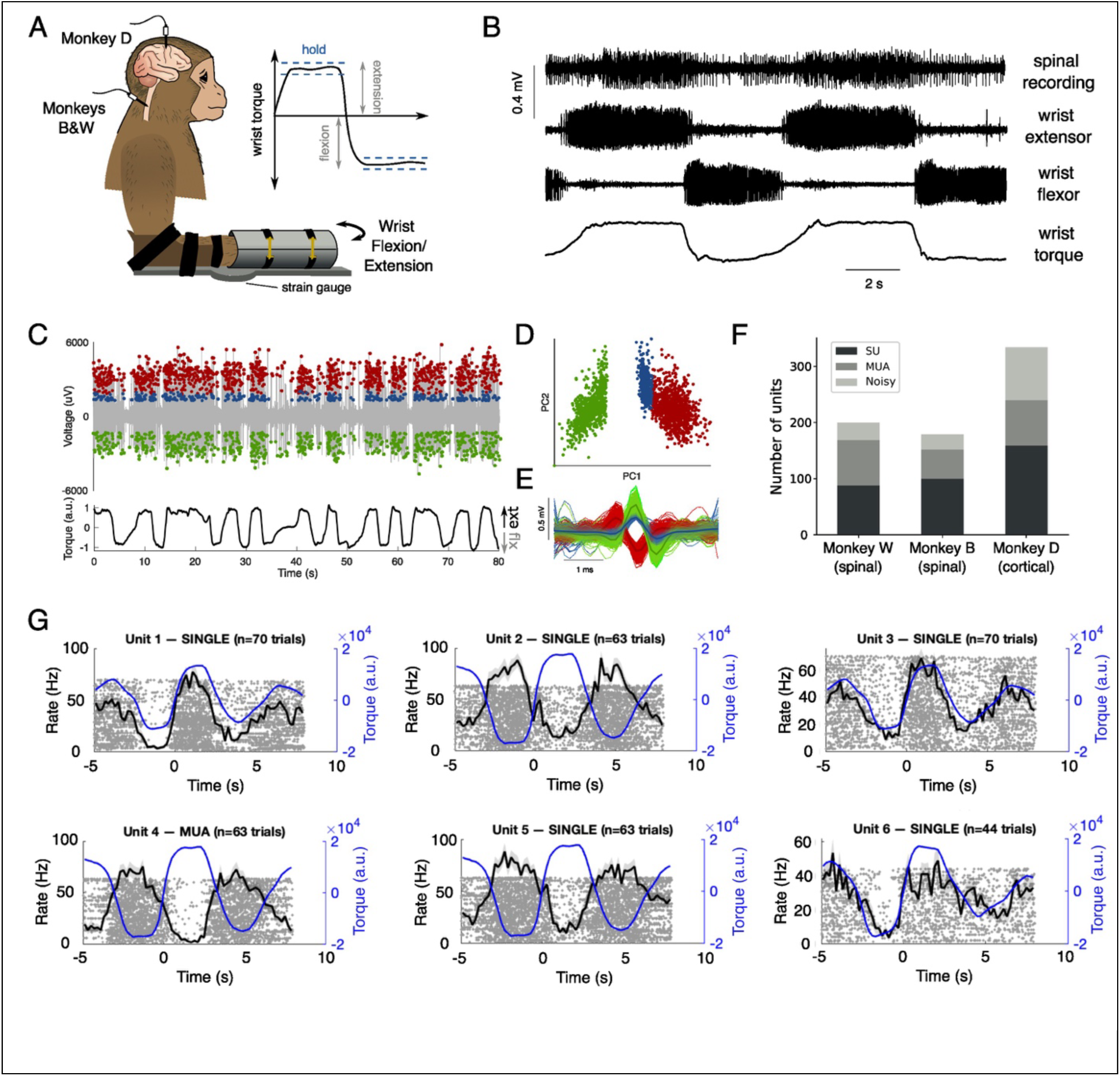
Cervical spinal recordings during isometric torque production. (A) Task schematic. Monkeys performed isometric wrist flexion or extension torque with visual feedback using a step-ramp-hold paradigm. Fingers were placed extended into a manipulandum equipped with strain gauges to record wrist torque. Spinal cord recordings (Monkeys B and W, cervical C6–T1) and motor cortex recordings (Monkey D, forearm area) were obtained during the same task. (B) Example raw traces from a spinal recording session showing simultaneous spinal cord extracellular recording, wrist flexor and extensor EMG, and torque signal during alternating flexion–extension trials. (C) Example spike sorting from a single cervical spinal electrode penetration showing color-coded sorted units overlaid on the voltage trace and concurrent torque signal. (D) Waveform clusters for sorted units in principal component space (PC1–PC2). (E) Average waveform shapes (± SD) for each sorted unit. (F) Unit counts across all three monkeys. Dark bars: single units (SU); medium bars: multi-unit activity (MUA); light bars: noisy. Monkey W (spinal): 88 SU, 81 MUA, 31 noisy. Monkey B (spinal): 100 SU, 52 MUA, 27 noisy. Monkey D (cortical): 159 SU, 81 MUA, 94 noisy. (G) Example spinal neuron peri-event time histograms showing directional tuning during the flexion - extension task. Six units shown (5 single units, 1 multi-unit). Black traces: trial-averaged firing rate. Blue traces: mean torque. Gray dots: spikes of individual trials (raster plot). Time aligned to torque reversal (time = 0).

Individual spinal interneurons showed clear directional tuning during torque production (Fig. 1G). The population divided into flexion-preferring, extension-preferring, and tonic units. Here, we focus on flexion- and extension-tuned units (see Extended Data Figure 6 for tonic units).

Despite robust single-neuron tuning, population-level analysis revealed no rotational dynamics in cervical spinal cord. We examined the structure of population activity by sorting units by their peak firing time and visualizing the resulting population firing rate matrix (Fig. 2A–C). Cortical units exhibited smoothly graded sequential activation, with firing rate peaks tiling continuously across movement phases (Fig. 2A). In contrast, spinal cord units showed abrupt, anti-phase modulation: the population split into two clusters that alternated between high and low activity states, with minor phase diversity (Fig. 2C). Electromyographic (EMG) activity of forearm muscles exhibited a similar but stronger binary alternating pattern (Fig. 2D).

**Figure 2.**
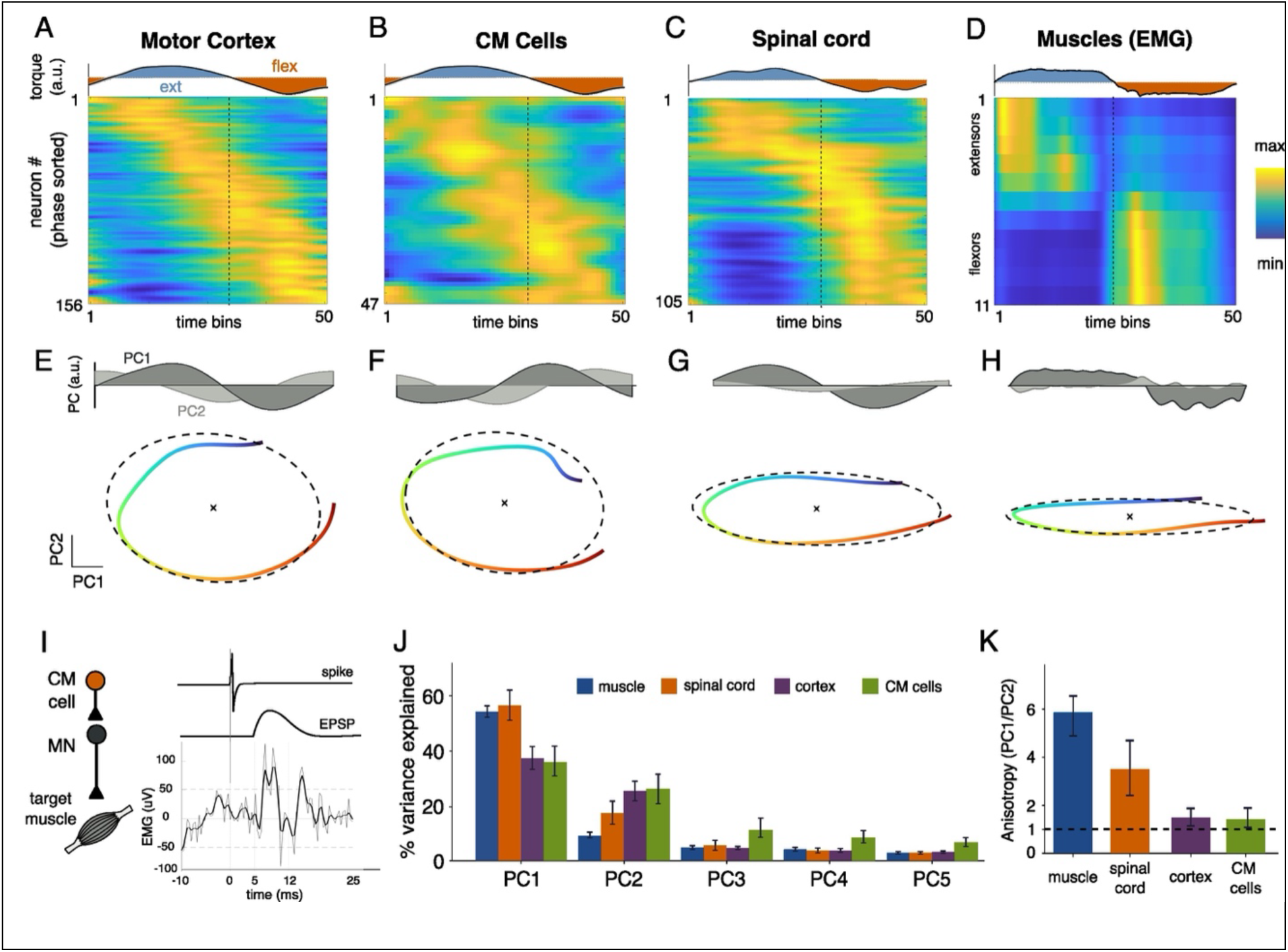
Distinct population dynamics across the motor neuraxis. (A–D) Population firing rate heatmaps for motor cortex (A; Monkey D, 156 task-modulated units; trial-averaged), CM cells from the same monkey (B; Monkey D, subset of 47 units; trial-averaged), spinal cord (C; Monkey B, 105 task-modulated units; trial-averaged), and forearm muscle EMG (D; 11 muscles, 6 flexors and 5 extensors; one example trial). Only units exceeding a modulation index threshold of 0.35 are included (the effect of varying modulation index is shown in Extended Data Figures). Units (and EMG) sorted by peak firing time. Cortical cells show smoothly graded sequential activation with continuous phase tiling. Spinal cord units show abrupt anti-phase modulation with two dominant clusters. Muscle activation shows a similar binary alternating pattern. Abscissa bins represent two sequential steps: first extension, then flexion. All trials were time-normalized to 50 time bins and time-aligned using torque reversal. (E–H) PC1 and PC2 vs time (top) and PCA trajectories in the PC1–PC2 plane (bottom) for motor cortex (E), CM cells (F), spinal cord (G), and muscles (H). Note that the PCA trajectories in H are for all recordings of Monkey B, not just the example trial shown in panel D. Solid lines indicate the trajectories (color-graded by movement phase; start in blue, end in red) and dashed lines indicate elliptical fits. Motor cortex trajectory is smooth, forming a close-to-circular loop; spinal cord and muscle trajectories are elongated along PC1 with minimal PC2 excursion forming ellipses. Each axis is scaled by the explained variance ratio of its PC, so that trajectories appear circular only when PC1 and PC2 carry comparable variance. While this scaling highlights the relative contributions of the two components, spinal trajectories remain elliptical and cortical trajectories remain near-circular without the weighting (see Extended Data Figure 5). (I) Example spike-triggered averaging effect from a CM cell onto a forearm muscle. Facilitation is observed in the 5-12 ms post-spike window. Dashed horizontal lines indicate the 95% CI of pre-spike EMG activity. Average EMG activity is shown with 1 SD shading. (J) Variance explained by the first five PCs for each recording site. Motor cortex and CM cells (purple and green, respectively) show distributed variance; spinal cord (orange, Monkey B) is dominated by PC1; muscle (EMG, blue) is strongly dominated by PC1. (H) Anisotropy ratio (PC1/PC2 variance explained) across recording sites. Values near 1 (dashed line) indicate multi-dimensional, rotational-like geometry; high values indicate one-dimensional alternating dynamics. Error bars represent 95% CI using bootstrap estimations.

Principal components analysis of rhythmic units (modulation index threshold of 0.35; see Methods for details) formalized this distinction. Cortical population trajectories traced smooth elliptical paths through the PC1–PC2 plane (Fig. 2E), with flexion and extension conditions following curved, looping paths characteristic of rotational dynamics^1,2^. Spinal cord trajectories were markedly different: they traversed elongated, nearly one-dimensional paths along PC1, with minimal excursion along PC2 (Fig. 2G). EMG trajectories showed a similarly collapsed geometry (Fig. 2H). We quantified this with the anisotropy ratio (PC1 variance divided by PC2 variance), where values near 1 indicate balanced, rotational geometry and large values indicate a single dominant dimension (Table 1, Fig. 2K). Cortical activity was close to balanced, consistent with two-dimensional rotational trajectories, whereas both spinal cord and muscle activity were strongly anisotropic, reflecting the dominance of a single flexion-extension alternating dimension. Critically, this difference in population activity dynamics between the motor cortex and the spinal cord was not driven by behavior: computed identically, the muscle PCA trajectories of all monkeys were strongly anisotropic and concentrated on PC1, unlike the cortical neural activity from those same animals (Table 1 and Extended Data Figure 9). While Monkey D performed shorter trials compared to the other two monkeys, resulting in larger PC2 explained variance (see Extended Data Figure 8 for details), torque PCA also resulted in PC1-dominated trajectories where PC1 accounted for most variance (Table 1). The elevated PC2 of cortical activity is therefore a property of cortical dynamics rather than of the movements the monkeys produced. See Extended Data Figures 1 and 3 for Monkey W PCA results and example units.

**Table 1.** PCA trajectory statistics for the neural population activity dynamics, muscle activity dynamics and torque profiles for all 3 monkeys.

| Monkey<br>(Region) | Data type | PC1 %explained<br>variance [95% CI] | PC2 %explained<br>variance [95%<br>CI] | Anisotropy ratio [95%<br>CI] |
| --- | --- | --- | --- | --- |
| Monkey D<br>(Motor Cortex) | Neural<br>population<br>activity<br>dynamics | 36 [33–42] | 26 [22–29] | 1.42 [1.14–1.86] |
| Monkey B<br>(Spinal Cord) |  | 56 [51–62] | 17 [13–22] | 3.29 [2.41–4.70] |
| Monkey W<br>(Spinal Cord) |  | 25 [22–31] | 11 [9–15] | 2.29 [1.65–3.10] |
| Monkey D | Muscle<br>activity<br>dynamics | 45.1 [41.2–49.1] | 8.9 [8.0–10.4] | 5.04 [3.94–6.00] |
| Monkey B |  | 54.6 [52.0–56.9] | 9.2 [8.5–10.6] | 5.93 [4.95–6.68] |
| Monkey W |  | 29.7 [27.5–32.3] | 7.4 [6.5–8.6] | 4.01 [3.37–4.96] |
| Monkey D | Torque<br>dynamics | 60.3 [59.2–61.6] | 10.3 [9.5–11.0] | 5.86 [5.41–6.36] |
| Monkey B |  | 78.8 [78.0–79.9] | 4.0 [3.5–4.7] | 19.75 [17.04–22.47] |
| Monkey W |  | 86.5 [85.4–87.8] | 2.3 [1.8–3.0] | 37.97 [29.08–48.89] |

To confirm that our spinal results reflect a genuine absence of rotational dynamics rather than methodological limitations, we demonstrate that motor cortex recordings from Monkey D (obtained in the same task, under similar behavioral performance, and using identical recording hardware, spike sorting pipeline, and analysis methods) showed clear looping trajectories (Fig. 2E), distributed variance across PCs (Fig. 2J), and a near-unity anisotropy ratio (Fig. 2K). This replicates previous cortical findings^1,2^ and confirms that our methods detect rotational structure when present. See Extended Data Figure 2 for example cortical units.

These findings reveal distinct computational dynamics at different levels of the motor system during the same voluntary muscle contractions. Spinal cord does not implement rotational dynamics during single-joint isometric torque control; instead, spinal population activity reflects rapid alternation between antagonist-related states, consistent with a direct transformation of descending commands into muscle activation patterns. The contrast between cortical and spinal population geometry (i.e. distributed and multi-dimensional versus concentrated and one-dimensional) suggests fundamentally different computational strategies: cortex generates flexible temporal sequences for movement planning and execution^2,3,17^, while spinal circuits implement rapid sensorimotor transformations^18,19^.

We also examined those cortical cells separately that had synaptic linkages to spinal motoneurons, i.e. corticomotoneuronal (CM) cells, as evidenced by post-spike facilitation/inhibition in spike-triggered averages of EMG (Fig. 2I). CM cell activity showed rotational dynamics similar to other cortical units, in difference to the dynamics of spinal neuron and forearm muscle activity (Fig. 2B,F). This indicates that CM cells, closely linked to spinal motoneurons and their innervated muscles, nevertheless shared activity dynamics with other cortical units as reflected in larger second principal components. This is coherent with CM cells’ activity being more influenced by their local input from layers II, III and VI, and from cortico-cortical projections than by indirect spinal feedback^20,21^.

Why do our findings differ from recent reports of spinal rotational dynamics^11–13^? A critical difference lies in task structure. Previous studies examined rhythmic multi-joint movements in which muscle groups are activated in stereotyped temporal sequences. During locomotion or scratching, multiple joints cycle through repeated phases, and neural populations tracking these patterns will necessarily produce trajectory rotation in state space -- not because spinal circuits perform computational rotations analogous to cortex, but as a geometric/biomechanical consequence of sequential activation across joints. This interpretation is supported by demonstrations that repetitive activation alone can produce rotational structure in population analyses^22^. Single-joint isometric torque production eliminates this confound; although torques alternated between extension and flexion these were separated by sustained static activation rather than involving rapid sequential cycling. Additionally, previous reports examined lumbar spinal activity during hindlimb movement in non-primates, whereas our recordings targeted cervical segments during primate forelimb control. While differences in species or segmental level may contribute, task structure provides the most parsimonious explanation.

Two possible limitations merit consideration. First, recordings were obtained from single-electrode penetrations across sessions rather than simultaneously. Second, the method used was blind to laminar identity of the recordings. However, motor cortex data collected with identical methods showed clear rotational geometry (Figure 2), indicating that non-simultaneous recordings (aligned to a behavioral marker) without laminar identity do not preclude detection of rotational dynamics. The consistency of results across both monkeys providing spinal recordings, in addition to the monkey used for cortical recordings, strengthens confidence that our findings reflect genuine differences in computational properties between spinal and cortical circuits.

We demonstrate that primate cervical spinal interneurons do not exhibit rotational population dynamics during alternating single-joint isometric contractions, in contrast with neurons of the motor cortex under identical task conditions. These results indicate that task structure critically influences population dynamics in spinal circuits and that rotational dynamics are not always a shared unique computational principle across the motor axis. Significantly, CM cells, i.e. corticospinal neurons with post-synaptic effects on motoneurons, had rotational dynamics similar to other cortical non-CM cells, rather than dynamics seen in the activity of spinal populations.

Together, this indicates that cortical circuits, during voluntary movements, accomplish functional control of fundamentally higher dimension than that of the spinal cord. This difference is directly explained by the distribution of phase preferences across each population: most recorded spinal units fired selectively during either flexion or extension, producing a bimodal population firing profile that is captured almost entirely by a single principal component, with little residual structure for a second component. Cortical units, in contrast, were tuned across a continuous range of movement phases, including intermediate phases between pure flexion and extension; capturing this distributed organization requires two principal components of comparable magnitude, yielding balanced PC1/PC2 amplitudes and a rotational trajectory in PCA space. Hence, these findings show that motor cortex and spinal cord activity exhibit distinctively different population dynamics during the same voluntary task.

Understanding how neural activity is transformed from cortical rotational dynamics to spinal alternating patterns and to muscle activation will be important for comprehending sensorimotor control, for determining the pathophysiology of motor disorders, as well as for developing neural prosthetics. For instance, understanding how the dynamics of motor cortex activity is transformed into spinal dynamics, and vice versa, would allow one to predict the spinal activity pattern evoked by a cortical stimulus (or the inverse: compute the cortical stimulus needed to evoke a target spinal pattern). In the context of spinal cord injury, for example, it might guide the design of brain computer interfaces that are coupled to stimulators below the lesion site^23–24^.

While many general features of motor control have been adequately described by optimal feedback control theory^25–26^, how these computations are neuronally implemented across the cortico-spinal (and other motor) systems remains poorly understood. Discerning the population and interaction dynamics of the cortico-spinal system provides key constraints to mapping out how control laws are implemented across the motor system, how they are affected in disease, and how neuromodulation might act on motor systems to restore effective upper limb control^27^.

## Online Methods

### Data collection

All procedures were approved by the University of Washington Institutional Animal Care and Use Committee. Neural data were collected from three adult rhesus macaques performing isometric wrist torque tasks (see Table 2 for summary). Animals were seated with the forearm secured in a manipulandum measuring torque about the wrist joint. Visual feedback displayed real-time torque as a cursor on screen. Monkeys B and W provided cervical spinal cord recordings; Monkey D provided motor cortex recordings. EMG recordings came from all three monkeys. All monkeys performed alternating step-ramp-hold torque trials (flexion or extension). Monkeys B and D performed trials to a single target torque level (1 s hold). Monkey W performed the same task with three randomly presented torque levels per direction. Trials were self-paced with variable inter-trial intervals (2–5 s). Extracellular recordings were obtained using glass-coated tungsten electrodes during separate sessions from cervical spinal cord (C6–T1, ipsilateral to the torque-producing arm) and the contralateral primary motor cortex (hand/forearm area). Intramuscular EMG recorded wrist flexor and extensor muscle activity (up to 16 muscles).

**Table 2.** Study subjects and behavioral tasks. Isometric tasks did not allow the wrist to move; auxotonic tasks allowed the wrist to move against a load (creating an equivalence between joint torque and joint angle). All tasks provided visual feedback based on wrist torque.

| Monkey | Sex | Tasks | Recording sites |
| --- | --- | --- | --- |
| Monkey B | M<br>(rhesus<br>macaque) | Single target step isometric<br>ramp and hold | Cervical cord gray matter<br>(C6-T1) |
| Monkey W | M<br>(pigtail<br>macaque) | 3 target step isometric ramp<br>and hold | Cervical cord gray matter<br>(C6-T1) |
| Monkey D | M<br>(rhesus<br>macaque) | Single target step isometric<br>and auxotonic ramp and hold<br>tasks | Motor cortex hand/forearm<br>area |

### Spike sorting and unit classification

Spikes were detected from raw neural recordings using a threshold of 2.5 standard deviations from baseline. Detected waveforms were sorted using principal components analysis followed by expectation-maximization clustering. Units were classified as noisy if ISI violation rate exceeded 5% or isolation distance was below 5. Among remaining units, single units were classified by meeting all of the following criteria: ISI violation rate ≤1%, isolation distance ≥20, and coefficient of variation of waveform amplitude ≤0.4. All other non-noisy units were classified as multi-unit activity. Since the maximum % ISI violation was <4% even in noisy units, units were classified as noisy based mostly on isolation distance and waveform stability (see Extended Data Figure 4). Therefore, noisy clusters were included in population dynamics analysis. Noisy clusters were only excluded when identifying CM cells.

### Population dynamics analysis

Spike trains were convolved with a Gaussian kernel (σ = 50 ms) to obtain continuous firing rate estimates and z-score normalized per neuron. For the population analysis, units with a modulation index (MI) below 0.35 were excluded to retain only task-modulated units, removing tonic units that do not differentiate between movement conditions. MI was defined as (*FR_max_* – *FR_min_*)/(*FR_max_* + *FR_min_*) where FR denotes firing rate. This yielded 105 units from Monkey B (spinal), 115 units from Monkey W (spinal), and 156 units from Monkey D (cortical). Varying MI did not qualitatively change the results; see Extended Data Figure 7 for more details. Trials were normalized to 50 time bins. For visualization, units were sorted by their peak firing time to construct population firing rate heatmaps (Fig. 2A–D). Principal components analysis was applied to phase-averaged population activity matrices (units × phase bins). To quantify the geometry of population trajectories, we computed the anisotropy ratio, defined as the ratio of variance explained by PC1 to variance explained by PC2. Values near 1 indicate roughly isotropic, multi-dimensional trajectories consistent with rotational dynamics; values substantially greater than 1 indicate elongated, one-dimensional trajectories consistent with alternating dynamics. EMG signals were rectified, smoothed, and analyzed similarly.

### Confidence interval estimation

Bootstrap confidence intervals (CI) were computed using resampled with replacement (n = 100 iterations); each iteration drew a full set of trials, computed the trial-averaged firing rates (or EMG profiles), and applied PCA to extract trajectory geometry metrics. The 2.5th and 97.5th percentiles of the resulting distribution defined the 95% CI for each metric.

### Data and code availability

Data and analysis code are available from the corresponding author upon reasonable request.

## Author Contributions

W.S.S., S.P., M.A.M., S.P., E.E.F. performed the recordings. J.B.N. digitized the original data from Fetz’s lab. O.R. and J.B.N. conceived the study. O.R. designed and performed all analyses and drafted the manuscript. J.B.N., E.E.F. provided supervision and edited the manuscript.

## Acknowledgments

O.R. used two LLMs (OpenAI ChatGPT 5.2 and Claude Opus 4.6/Sonnet 4.5) to improve the readability of the manuscript. All outputs of LLMs were thoroughly revised and edited, and we assume full responsibility for the scientific integrity and accuracy of the manuscript. O.R. is supported by a Lundbeck Foundation Postdoctoral Fellowship (Grant no. R483-2024-1882).

---

**Extended Data Figure 1.**
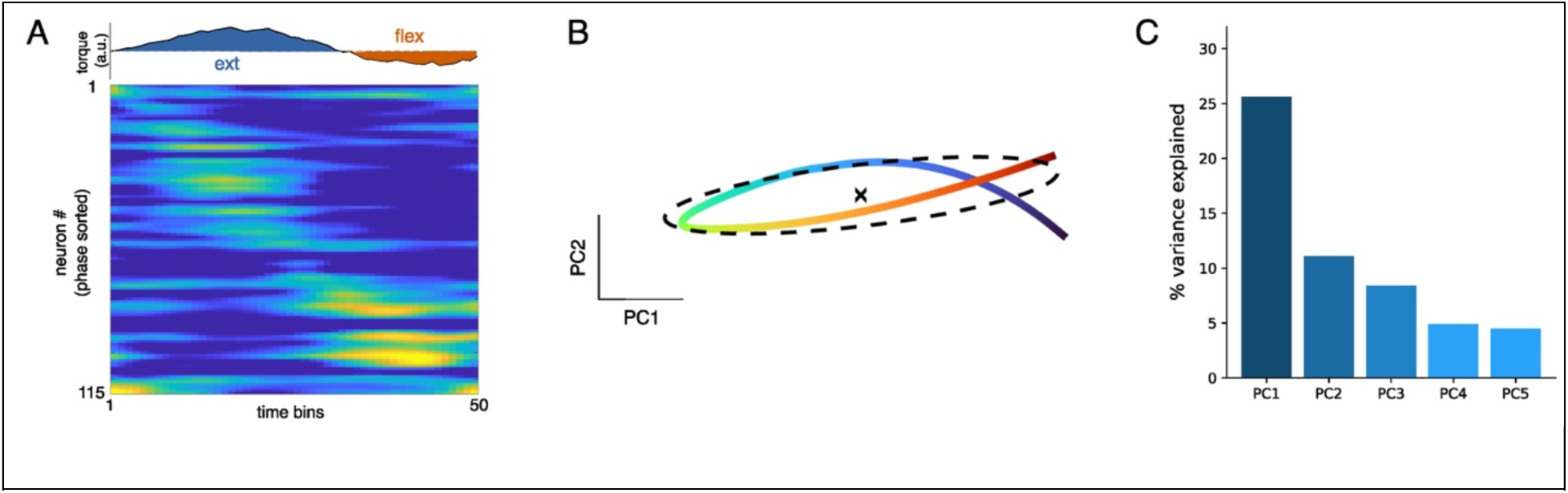
Monkey W PCA results for spinal populations. A. Neuron firing rate heatmap (MI threshold = 0.4). Two clusters at flexion and extension are visible (middle cluster, versus top and bottom). B. PCA in 2D of the heatmap shown in A. PCA is weighted by variance explained. Dashed line shows elliptic fit (aspect ratio = 6.4). C. Variance explained is dominated by the first principal component as observed in Monkey B.

**Extended Data Figure 2.**
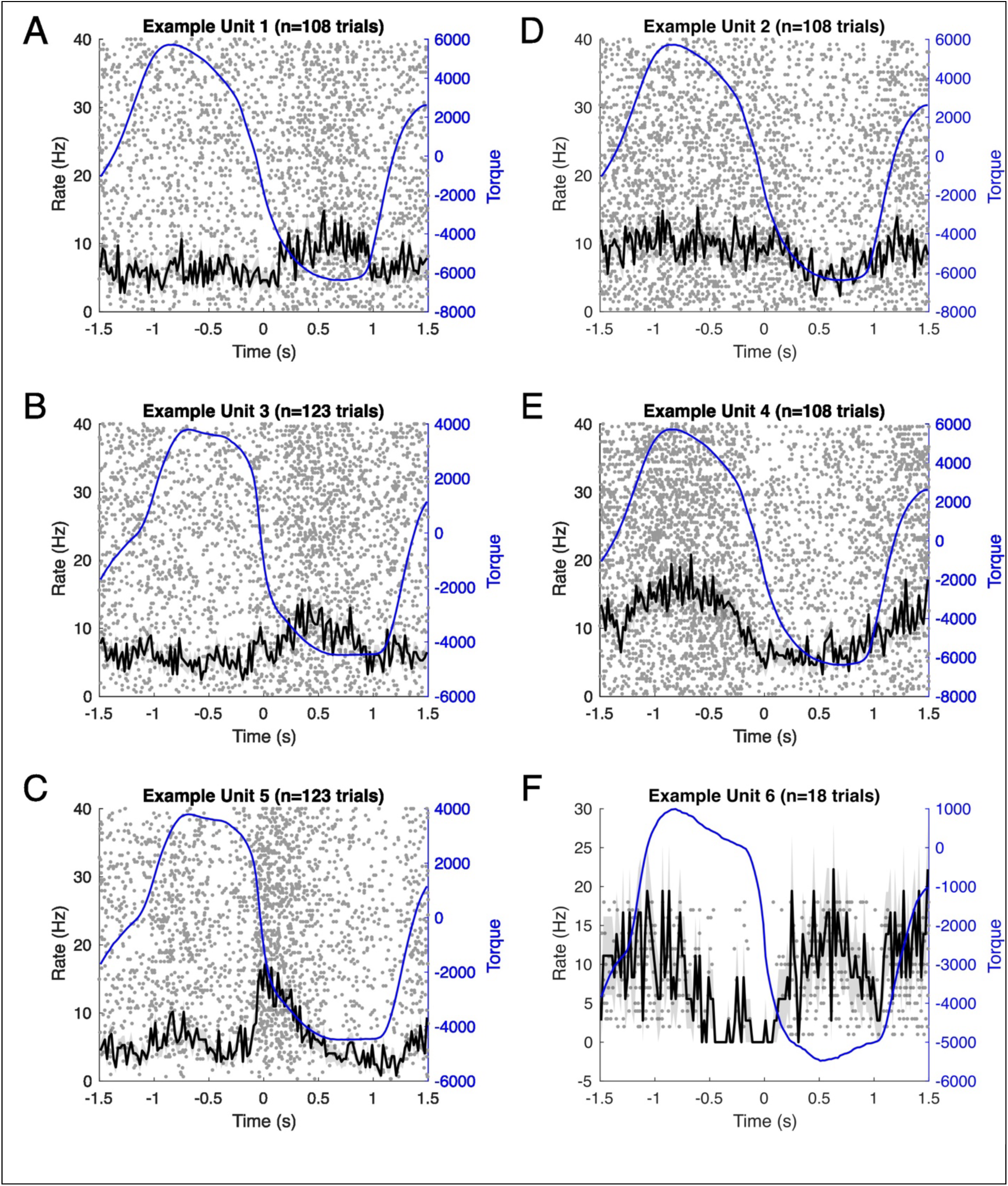
Example cortical units from Monkey D. Note the progression of phase offset from torque in panels A-F. Cortical units were phase locked to all phases of movement, unlike spinal units that were faithfully phase locked to flexion only, extension only, both (observed in Monkey W) or were tonic. Cortical units also showed more complex profiles (not shown here).

**Extended Data Figure 3.**
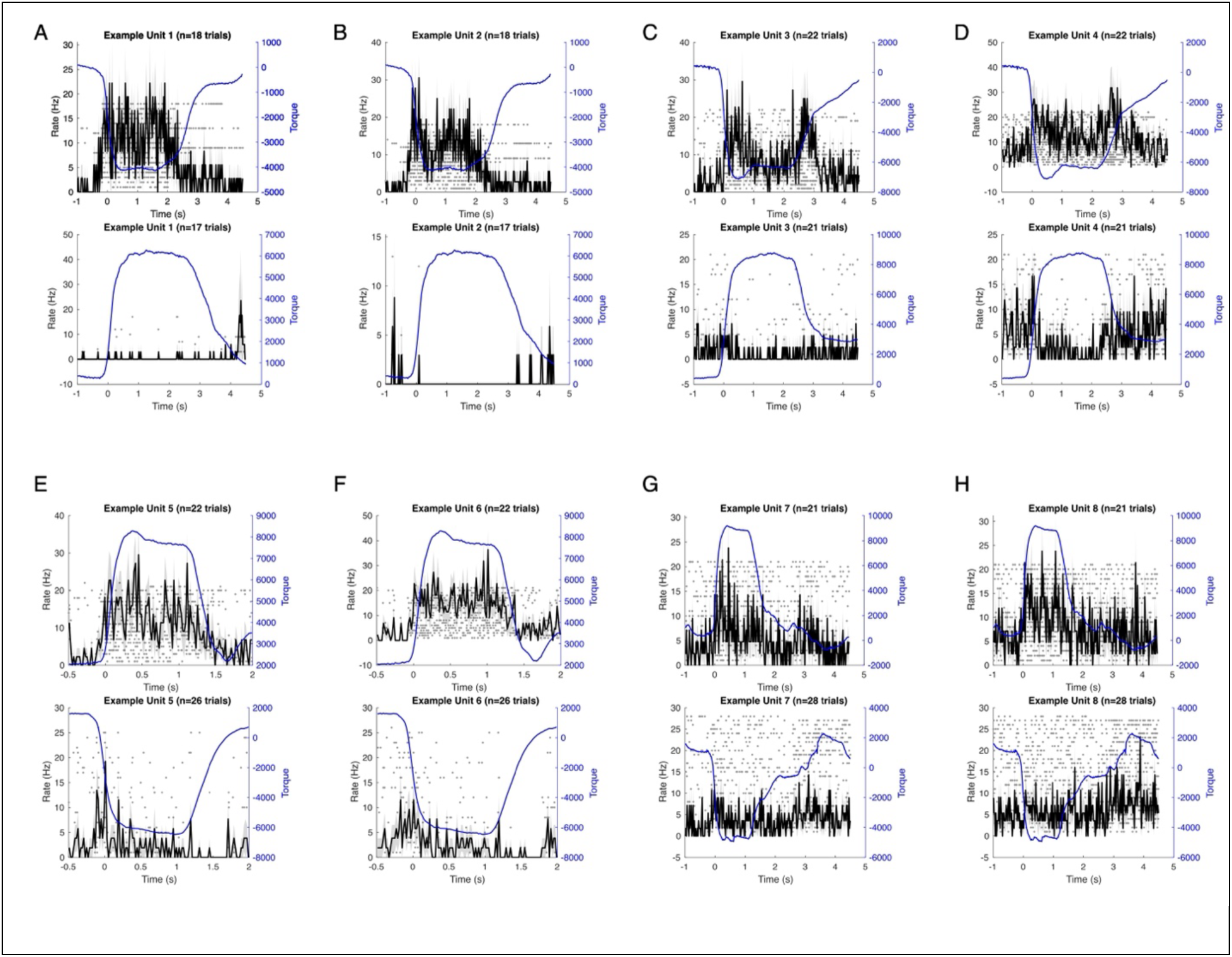
Eight example spinal units from Monkey W. A-D. Four units tuned to flexion. PSTH from the same unit during flexion trials are shown in the top row, and during extension trials in the bottom row. Notice that units are silent during extension, and active during flexion. E-H. Four spinal units from Monkey W tuned to extension. Some units were observed in Monkey W that were active in both flexion and extension (not shown here).

**Extended Data Figure 4.**
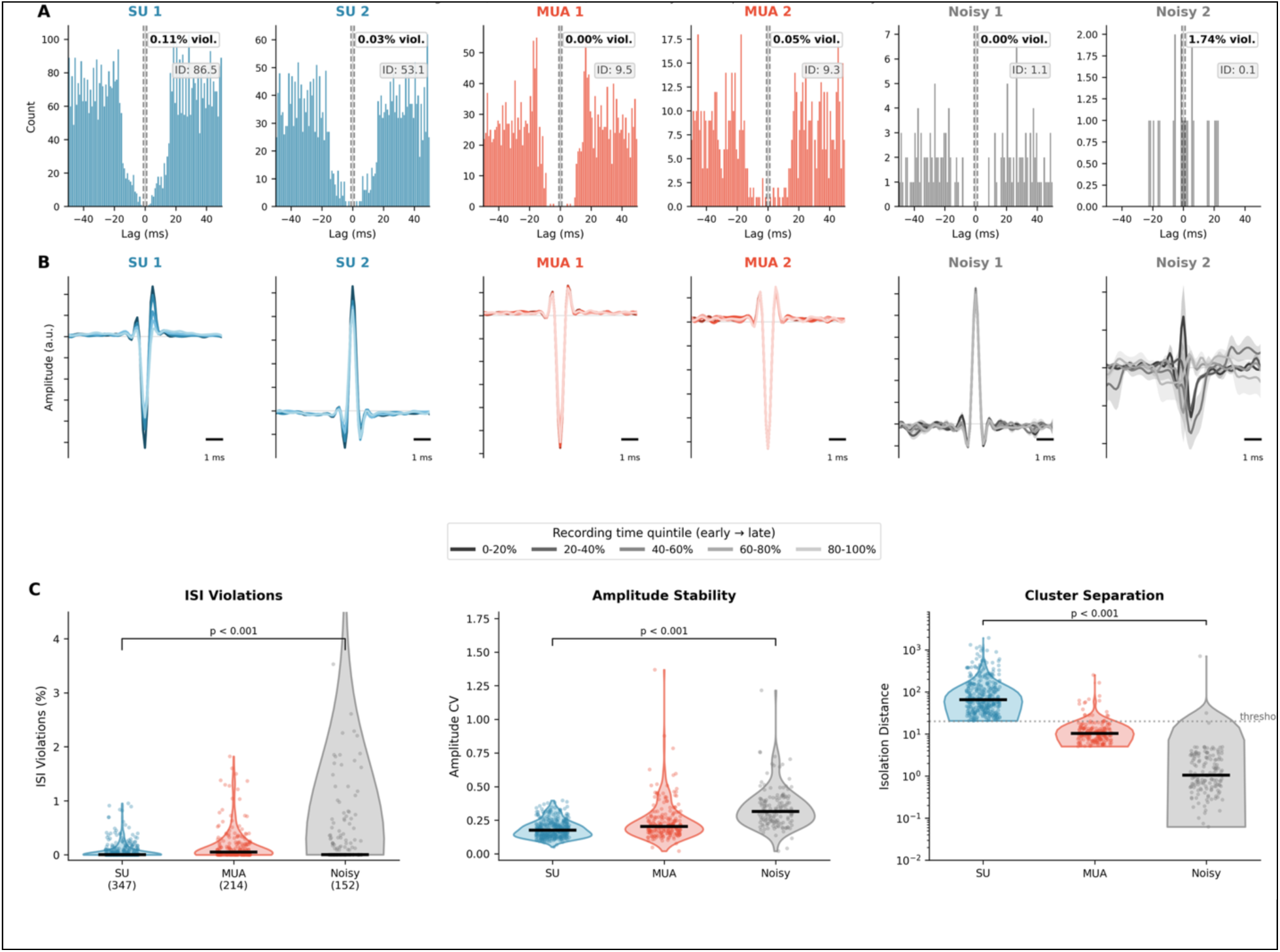
Unit quality across monkeys. Data for 347 single units (SUs) and 214 multi-unit activities (MUAs) and 152 noisy clusters (Noisy) pooled across all 3 monkeys. A. ISI histograms for 2 SUAs and 2 MUAs and 2 Noisy. Percentage violations and isolation distances (ID) are annotated on the histograms. Notice how the ISI violations are rare for all cluster types; units are, in effect, mostly separated by their amplitude stability and isolation distance. B. Stability of waveform throughout the recording. Each waveform is the average of all waveforms in these 5 intervals. Noisy clusters also often showed stable amplitudes throughout the recording, but had poor cluster separation compared to SUs and MUAs. C. % ISI violations, amplitude stability and cluster separation (isolation distance) comparison for all SUs, MUAs and Noisy clusters pooled across all monkeys. ISI violations were <4% for all cluster types, and hence waveform amplitude stability and cluster isolation distance were mostly the determining factor for classifying a unit as Noisy. Since quality of Noisy clusters showed acceptable ISI violations and amplitude stability, they were included in population dynamics analysis, but not in CM cell identification. Comparisons are using Mann-Whitney U-test, and p-values are corrected for multiple comparisons using the Benjamini-Hochberg method for false discovery rate.

**Extended Data Figure 5.**
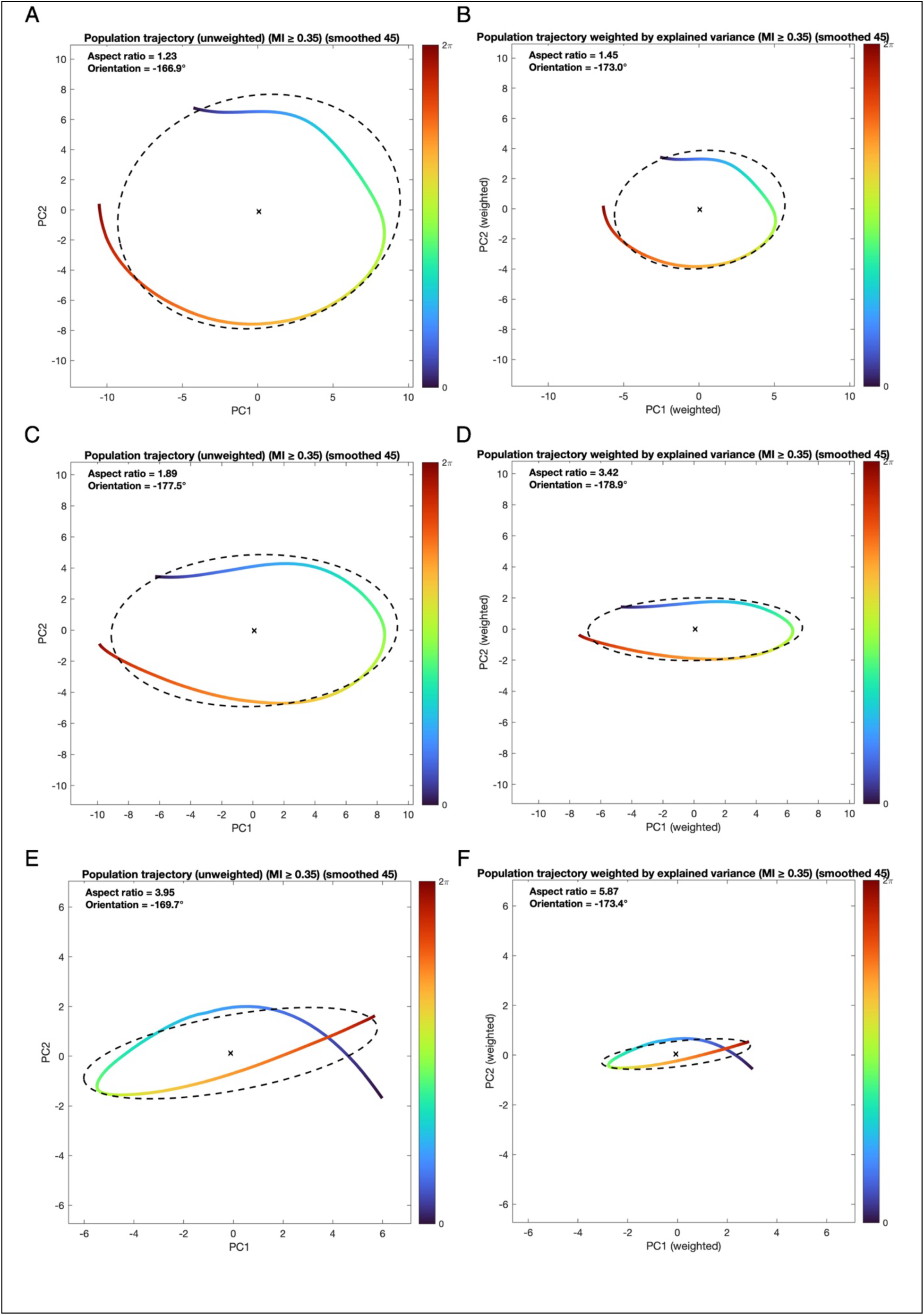
Unweighted versus weighted PCA trajectories. Rotational population dynamics should show close-to-circular trajectories in PCA space even after weighing the trajectories by explained variance of the respective components. Population dynamics with heavy alternating components should look more elliptical after weighing. A-B Monkey D. Weighting PCA trajectories by variance explained by each PC minimally affects the aspect ratio of the trajectory (compare to unweighted in panel A). Weighting trajectories by explained variance significantly increases the aspect ratio in both Monkey B (C-D) and Monkey W (E-F) because the dynamics of their spinal interneuronal activity were dominated by PC1, whereas that of Monkey D (cortical) was balanced between PC1 and PC2.

**Extended Data Figure 6.**
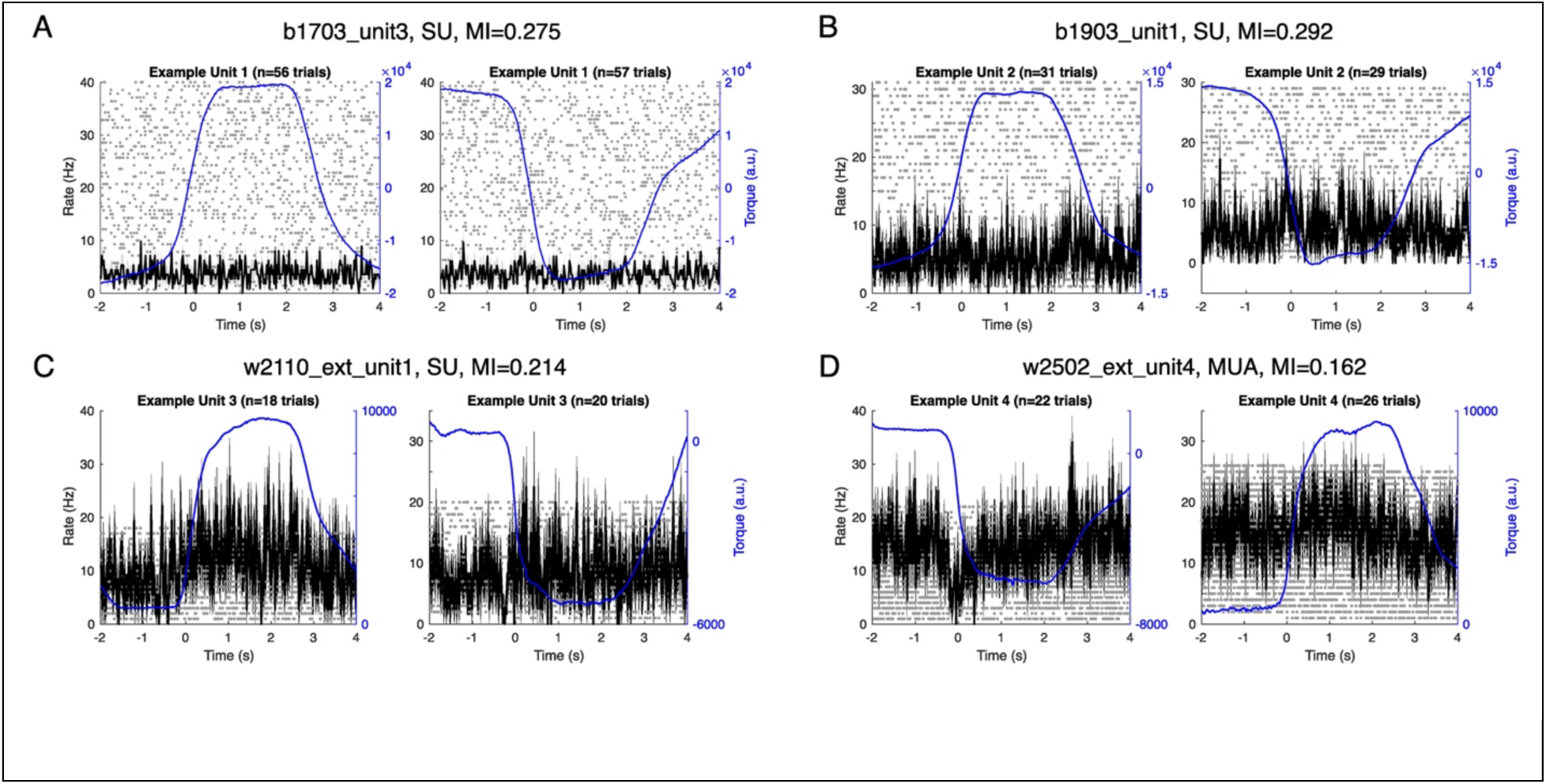
Example spinal units with low depth of modulation that are tonic or nearly tonic. A-B are PSTHs of two example units from Monkey B, each centered around flexion and extension trials. C-D shows two units from Monkey W.

**Extended Data Figure 7.**
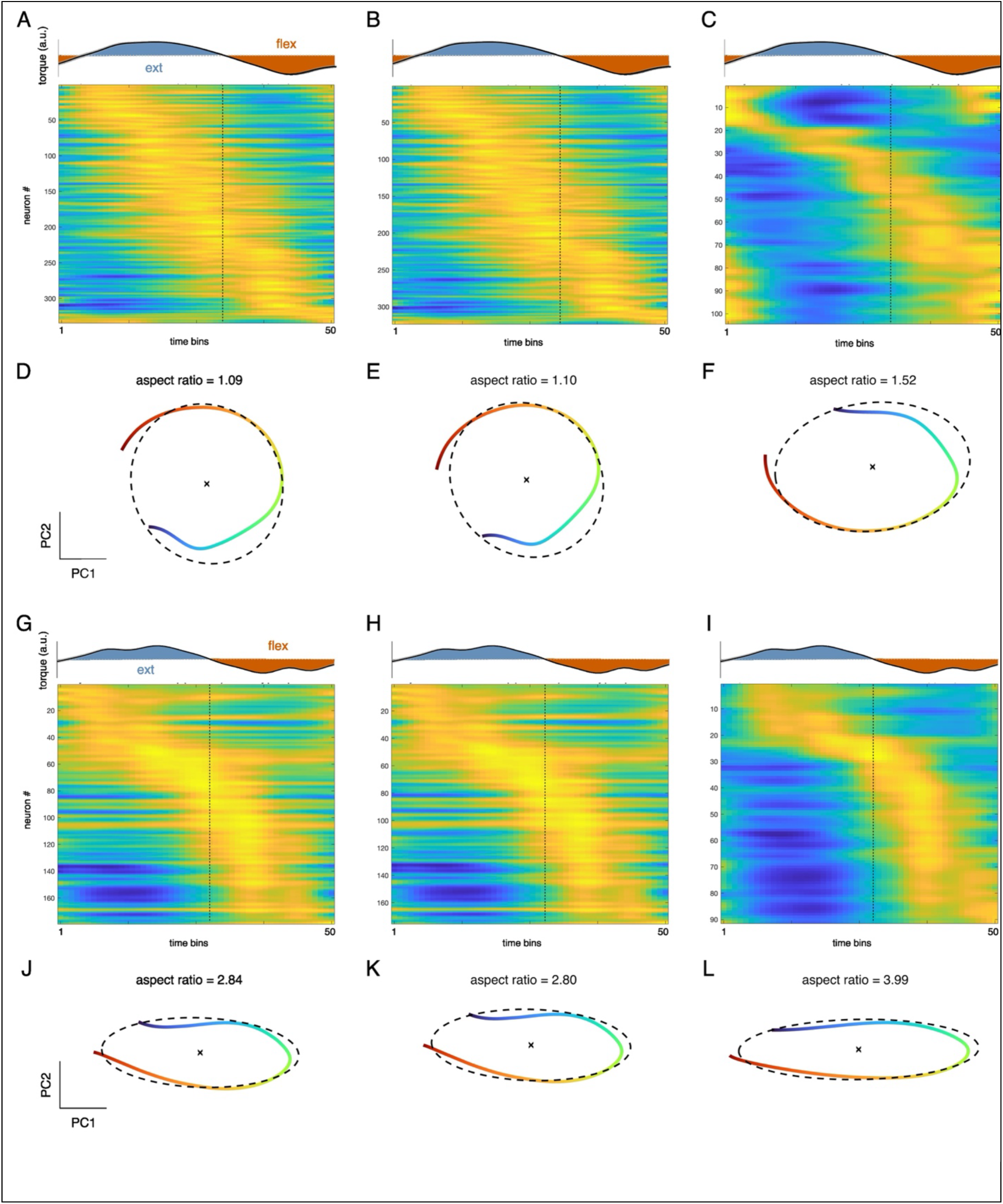
Unit firing heat maps and PCA trajectories with progressively increasing modulation index (MI) threshold. A-C: Monkey D with MI = 0, 0.2 and 0.4, respectively. D-F Corresponding PC1-PC2 trajectories corresponding to panels A-C. Trajectories are weighted by explained variance (see Methods). Rotational dynamics is maintained with progressively increasing MI threshold. G-L: Same panels for Monkey B (spinal). Increasing MI mostly removes tonic units. Alternating dynamics persist even at MI threshold = 0.

**Extended Data Figure 8.**
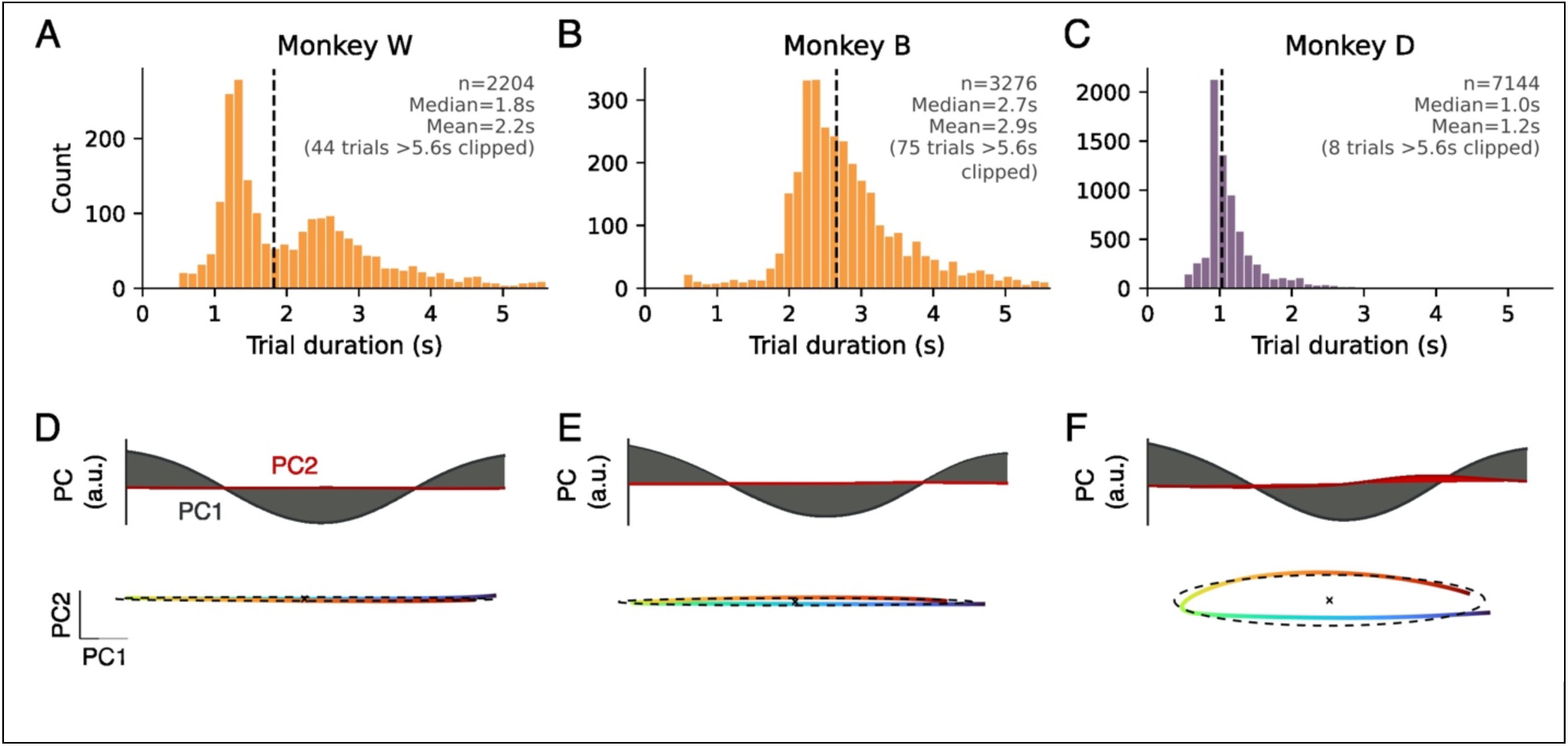
Task differences between monkeys. A-C Distributions of trial durations for each of the monkeys. Note that Monkey D had shorter trials than both spinal monkeys (albeit Monkey W has a large cluster of similar trial lengths). D-F Weighted PCA trajectories of time-normalized torque profiles (the same method of computing firing rate PCA trajectories) across all recordings for all 3 monkeys. The shorter trials correspond to the ramp phase being a larger proportion of the trial (i.e. shorter holds), which is reflected in a larger second principal component for Monkey D compared to Monkeys B and W. The torque profiles, however, are highly one-dimensional in PCA space, which indicates that task differences between the two spinal and the one cortical monkey are not sufficient to explain the differences in neural population dynamics reported.

**Extended Data Figure 9.**
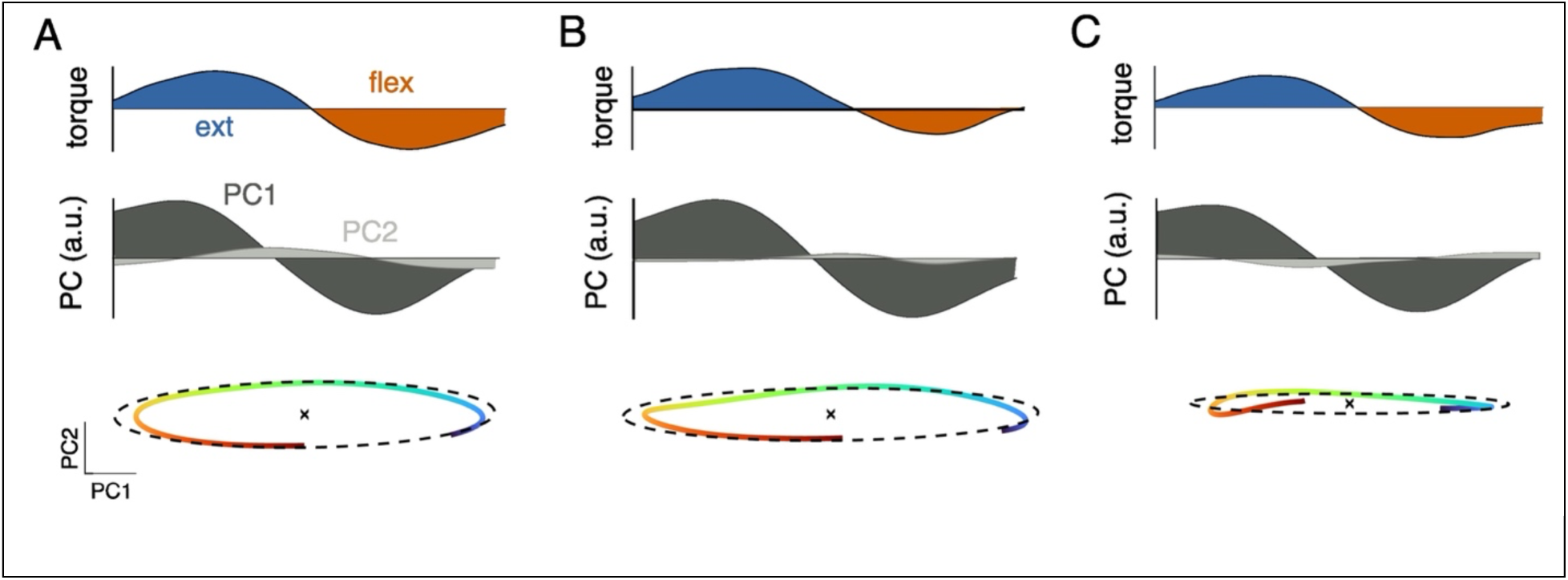
Weighted PCA trajectories of EMG activity for all 3 monkeys. A-C Monkey D, B and W, respectively. Notice that trajectories were dominated by PC1 for all 3 monkeys, indicating that differences between cortical and spinal neural population activity dynamics were not driven by task/motor output differences.

